## Supplemental Material for "Identifying Reproducible Transcription Regulator Coexpression Patterns with Single Cell Transcriptomics"

***Supplemental Material table of contents***

*Commentary on data representation ……………..………………………………….….........………2*

*Negative coexpression and candidate repression ………………………………….….........………4*

*Aggregate ribosomal profiles …………………….…………..………………………….........…….…8*

*ChIP-seq scoring and analysis …………………….…………………………………….........………8*

*Literature curation benchmark …………………….…………………………………….........…….…9*

*Supplemental figures ……………………………………………………………………….…..…….. 12*

*Supplemental citations ………………………………………………………………………..…..….. 20*

**Commentary on data representation**

This study focuses on identifying replicable single-cell coexpression patterns across a large corpus of data, setting aside questions of context-specific patterns (e.g., cell type, disease state, developmental stage) to first establish baseline expectations for generalizable signals. At the time of writing, we utilized all available data from the CellXGene resource, supplemented with data from GEO while taking care to include large tissue atlas studies. However, this collection still contains appreciable biological and technical variability and represents only a subset of all available scRNA-seq data. Recognizing these limitations, we aimed to identify potential sources of bias in our estimates of global TR coexpression profiles.

We first examined technology differences, noting that most datasets were generated using 10X Chromium strategies, but some used other platforms (Figure 1A; Supplemental Table 1). A useful comparison came from GSE132042, which included both 10X 3’ v2 and Smart-seq2 data from the same mice and matched cell types, minimizing inter-group variability. Using our aggregation approach, we calculated coexpression separately for each platform and compared the Top_200_ coexpression partners for each TR. Cell counts per cell type differed significantly between platforms, with 10X data often having an order of magnitude more cells. We chose not to subsample to equalize counts, as the inherent differences in cell count versus sequencing depth are a defining feature of 10X and Smart-seq platforms, making subsampling an unfair comparison

In Supplemental Fig. 2G, we plot the distribution of Top_200_ values for the 1,346 TRs that were measured in either the 10X or Smart-seq2 dataset, along with a null distribution from random sampling. While the overlap is significantly better than random (Wilcoxon test p.value < 2.2e-16), it is small with a median value of 10/200. We further note that the most similar (by Top_200_) TR profiles between technologies were aligned with the most consistent TRs presented in Fig. 2, led by Mxd3 (124/200), Foxm1 (114/200), and E2f8 (103/200). This outcome aligns with our prior observations of variability when comparing scRNA-seq datasets with similar designs, such as disease states. These findings reinforce the motivation for this study: to determine what patterns remain consistent across a large, diverse dataset collection.

Next, we created a metric to score how well each dataset agreed with the global aggregate, aiming to identify trends such as technical factors or tissue overrepresentation. For each dataset, we compared the Top_200_ overlap of its TR profiles against the global TR profiles (as shown in Figs. 2C,D, left panels). These overlaps were rank standardized, with a value of 1 indicating the best dataset match to the global profile for a given TR and 0 indicating the lowest overlap. A single dataset could score 1 for multiple TRs. The values were then averaged into a dataset-level vector, termed *Global agreement*, where higher scores reflect stronger alignment of a dataset’s TR profiles with the global profile. Results for human datasets are shown in Supplemental Fig. 2H.

In human, the dataset with the highest *Global agreement* was a large study of diseased and healthy gastrointestinal tissues (NowickiOsuch2023; CellXGene link: <https://cellxgene.cziscience.com/collections/a18474f4-ff1e-4864-af69-270b956cee5b>). In contrast, the lowest score belonged to a pancreas study that assayed a small number of cells on a rarely used platform (GSE85241; <https://cellxgene.cziscience.com/collections/6e8c5415-302c-492a-a5f9-f29c57ff18fb>). Notably, tissue type did not appear to drive high scores; for example, another intestinal dataset was among the lowest scores (Wang2020; <https://cellxgene.cziscience.com/collections/ff668d5d-5b3f-49ee-a007-ff0664bf35ec>). Instead, dataset size and platform, particularly 10X, were the most apparent factors (Supplemental Figs. 2I-K).

The difference in representation in the corpus between 10X and other methods reflects the current state of the field. Most scRNA-seq data, including large tissue atlas studies that are more likely to align with global patterns we seek to identify, are generated using 10X technology. Importantly, we still observe non-10X experiments with high *Global agreement*, and, given that we do see a (modest) agreement in the TR profiles generated between technologies (shown in the GSE132042 comparison), any common patterns will still be emphasized by our aggregation approach.

**Negative coexpression and candidate repression**

In this manuscript we predominantly focus on positive coexpression. As demonstrated by in Fig. 1D-F, the most reproducible positive coexpression patterns tend to be larger in magnitude and more consistent than the equivalent extremes for negative coexpression. We have observed this trend before in bulk coexpression metanalysis (Lee et al., 2004) and it has also been noted in work motivating the popular SCENIC workflow for single cell data (Aibar et al., 2017; Van De Sande et al., 2020). In addition, we found that the null overlap comparisons (looking at the intersect of the most positive/negative coexpressed gene partners of shuffled TR profiles) consistently gave higher median values for the Bottom*_K_* nulls than the Top*_K_* nulls (Figs. 2A,B; Supplemental Figs. 2A,D), which may be a consequence of our aggregation framework. Collectively, we have less confidence that the observed reproducibility of negative coexpression is driven by dynamic, biological regulatory processes rather than effects of low expression or technical noise.

Moreover, the focus of this paper is ultimately to nominate candidate gene targets of transcription regulators (TRs). Biochemically demonstrated regulatory interactions (the principal source of independent validation used in this paper) are heavily skewed towards activation (Chu et al., 2021), which focused our priorities. There are additional complications of using gene coexpression to nominate repression compared to activation. A predicted positive interaction means that the transcripts of a TR and candidate target are both detected across samples. A negative coexpression meanwhile can result when either the TR is well-expressed and a candidate lowly expressed, or inversely when the candidate is well expressed and the TR lowly expressed.

The scenario of a gene being well-expressed only in the absence of a TR is sensible in the context of repression, but it necessitates arguing that the absence of TR transcripts is causal in the abundance of candidate gene transcripts. This is inherently a weaker line of evidence than observing co-abundance. In the case of a TR being well-expressed and repressing a candidate, it is possible that the repression results in the complete cessation of the target’s expression, meaning that there is no variation in which to calculate coexpression. In sum, nominating repression using transcriptomics alone is arguably best handled by a differential expression framework that ensures that the TR is expressed.

Acknowledging these caveats, we still sought out to explore candidate repressive interactions, or TRs that may be described as more repressive in nature, as observed by their coexpression patterns.

We first examined the degree of overlap of a TR’s negative coexpression profile (Bottom_200_) across studies. The maximal values were slightly depressed relative to those from the Top_200_ comparisons. The largest human Bottom_200_ (158/200) belonged to the repressive nuclear receptor *NCOR1* in a comparison between profiles generated from tissue atlas resources (Uhlén et al., 2015; The Tabula Sapiens Consortium et al., 2022). In mouse, the largest overlap (Bottom_200_ = 149/200) corresponded to *Elk3* between studies of the blood-brain barrier (van Lengerich et al,. 2023) and aging brains (Kaya et al., 2022). Consistent with the Top_200_ comparison, the globally most similar data pairs were often derived from studies examining similar biological contexts.

As with the positive coexpression analysis, we averaged these pairwise comparisons to describe how consistent a TR is in its negative coexpression profile (Supplemental. Fig. 2). Unlike the mean Top_200_ comparison, most TRs did not have a mean Bottom_200_ that was greater than all the means generated across 1000 shuffled nulls, although an appreciable proportion did (human 28%, mouse 23%). Examples of regulators with consistent negative coexpression profiles in both species include the aforementioned *NCOR1*, the histone methyltransferase-encoding *KMT2C*, nuclear factor *NFAT5*, and the dual RNA and DNA-binding *SON*.

In an attempt to identify candidate repressive factors, we examined whether any TRs had more consistent negative correlation profiles than positive correlation profiles, relative to the respective nulls (Figs. 2A,B; Supplemental Figs. 3C,D). We subset TRs to those with an average *Top_K_* value that was less than the typical null *Top_K_* value, indicating TRs with weak cross-dataset agreement in their positive coexpression. We then plotted the mean *Bottom_K_* values of these TRs against the mean *Bottom_K_* values across all 1000 null iterations, to see if any of these subset TRs had a mean Bottom_200_ value appreciably different than the range of nulls.

Using *K*=200 revealed few convincing examples of TRs whose negative, but not positive, coexpression was consistent across datasets (Supplemental Figs. 3C,D), That is, most TRs that had weak consistency in their positive coexpression also had weak consistency in their negative coexpression. Relatedly, TRs with the most consistent Bottom_200_ values also tended to have Top_200_ values exceeding the null expectation and were thus excluded from this analysis. Using *K*=1000 did however provide examples of TRs whose negative, but not positive, coexpression profiles exhibited greater overlap than shuffled profiles. Notably, zinc finger TRs were well represented in this analysis. A leading example in human was the Ikaros zinc finger *IKZF5,* a hematopoietic TR implicated in platelet deficiency, and which is reported to be understudied relative to other Ikaros genes (Lentaigne et al., 2019).

Regarding the cross-species comparison, Fig. 4A demonstrates that the Bottom_200_ overlap for most orthologous TRs was greater than the typical shuffled null value. We also repeated the ortholog retrieval score represented in Fig. 4B (which focuses on positive coexpression) using Bottom_200_ values. To reiterate, this analysis queries one TR’s most consistently coexpressed partners against each TR in the other species and returns the quantile (from 0 to 1) of its matched ortholog. A value of 1 means that a TR had more overlap with its ortholog than with any TR in the other species, indicating specificity in its shared coexpression partners.

We show the result of this analysis in Supplemental Fig. 6C, which demonstrates a large degree of specificity in the preservation of negative coexpression partners between orthologous TRs, albeit attenuated compared to the reproducible positive interactions (median *Bottom_200_ human in mouse* = 0.88; *mouse in human* = 0.93). 22 TRs achieved a perfect *Bottom_200_* ortholog retrieval score for both species, with ZNF532 (mouse Zfp532) having the largest overlap of this set (*Bottom_200_* = 107). This TR is not well-represented in the literature, and thus it is an open question as to why this zinc finger regulator is exceptional in its consistency of shared negative coexpression partners across species.

Each TR can be represented by a pair of *human in mouse* and *mouse* *in human* ortholog retrieval score separately for its positive and negative profiles. To summarize this information, we averaged each pair (i.e., *Bottom_200_ human in mouse* and *Bottom_200_ mouse in human*), and plotted these values against each other in Supplemental Fig. 6D. This revealed numerous examples of TRs that shared more negatively and positively coexpressed partners with their ortholog relative to other TRs. For example, in the main text we note that NEUROD6 had a perfect *Top_200_* ortholog retrieval score for each species (and thus its average is also 1), and its averaged *Bottom_200_* ortholog retrieval score was 0.99. We caution that the size of this negative coexpression overlap is small relative to the positive overlap (*Top_200_* = 50/200, *Bottom_200_* = 9/200), but the high *Bottom_200_* ortholog retrieval score still indicates that this overlap is greater than with the vast majority of other TRs. Notably, four TRs — SOX17, SOX18, SUB1, and THRA — achieved a perfect ortholog retrieval score across all comparisons, which suggests that these TRs are exceptionally conserved in their most extreme coexpression partners.

Lastly, we re-evaluated the coexpression benchmark by reversing the ranks to prioritize negative coexpression for scoring. Our goal was to identify any TRs that performed poorly using positive coexpression but displayed improved performance using negative coexpression, with the aim of uncovering repressive interactions. Prioritizing negative coexpression greatly diminished performance for most TRs relative to positive coexpression and binding (Supplemental Fig. 5B), consistent with the observation that the literature corpus is enriched for activating interactions (Chu et al., 2021). We did, however, identify 25 TRs in human and 23 in mouse that exhibited enhanced performance, with 9 of these factors occurring in both species. Moreover, 12 of these 25 human TRs and 8 of the 23 mouse TRs also had performant binding aggregations (*Quant_binding* > 0.9), and three were common to both species: MAX, NRF1, and YY1. Collectively, these observations may point to TRs with more readily identifiable repressive activity: their negative, but not positive, coexpression profiles are able to retrieve their curated targets relative to sampled targets, and these curated targets are also associated with elevated binding signal in the respective TR ChIP-seq experiments.

**Aggregate ribosomal profiles**

In keeping with our use of ribosomal genes as a positive control for known coexpression, we generated aggregate profiles for each of the 82 highly conserved L/S ribosomal genes. We then examined their top 82 ranked coexpressed gene partners to validate that the top of their aggregate profiles was enriched for other ribosomal genes (Supplemental Fig. 4A,B).

In both mouse and human, we found that, on median, 71 out of the 82 top ranked partners were other L/S ribosomal genes. Note that these values are slightly depressed by our choice to limit the analysis to one-to-one orthologs; in most cases, additional ribosomal genes are in the top 82. This validates the prioritization of known biological coexpression. The notable exceptions fortuitously illustrate a point about tissue-specific patterns. One was human *RPL39L*, a paralog of *RPL39,* which has been described for its involvement in alternative ribosomal activity in the testes and spermatogenesis (Li et al., 2022). We observed that the top ranked *RPL39L* coexpressed partner was *PBK*, also implicated in testes function and spermatogenesis (Miki et al., 2020). Similarly, *RPL3L*, a heart and skeletal muscle-specific paralog of *RPL3* (Shiraishi et al., 2023) had only 2 out of 82 of its top partners belonging to the L/S ribosomal set. Notably, the cardiac myosin genes *MYH7* and *MYL2* ranked 5th and 6th, respectively, in *RPL3L’s* aggregate ranking of coexpressed partners. These results show that while robust context-independent coexpression patterns can be readily observed, context-specific patterns can also be discovered in our data.

**ChIP-seq scoring and analysis**

Each Unibind ChIP-seq experiment is represented as a table of genomic ranges representing the sites where the canonical TR motif within an inferred binding event (“peak”) was identified. GenomicRanges (Lawrence et al., 2013; V1.50.2) was used for all analyses. For each experiment we scored gene binding as in prior work (Morin et al., 2023), using a continuous scoring function based on the exponential decay function introduced by Ouyang et al. (2009):

$$S_{g}= \sum_{k=1}^{K} e^{-\frac{d_{k}}{d_{0}}}$$

Where *S* is the binding score for a gene (*g*) in one TR experiment, *K* is the number of peak summits within 1Mbp of the gene TSS, *d_k_* represents the absolute distance in bps between the TSS and the peak summit, and *d_0_* is the decay constant, set to 5,000 as in the original publication.

The same ChIP-seq experiment may be represented multiple times within Unibind, corresponding to TRs matched to distinct canonical motifs. In such cases, we averaged the duplicated binding scores such that each experiment was only represented once. To alleviate batch/technical considerations, we bound all de-duplicated gene binding vectors into a gene by experiment matrix, added 1, and applied a log_10_ transformation followed by quantile normalization (preprocessCore R package version 1.48). This process was done separately for each species' experiments. To generate an aggregate binding profile, we averaged the gene binding vectors specific to each TR.

Regarding Figure 6, ASCL1 had duplicated ChIP-seq datasets in Unibind and so we took the union of peaks for datasets with the same experiment ID. All peaks were resized to ~300 base pairs, and a “consensus” list of bound regions was generated as a union of all discrete bound regions across datasets. We calculated the count of individual datasets that overlapped this consensus set and plotted these counts using igvR (Shannon 2023; V.1.22).

**Literature curation benchmark**

A primary challenge in gene regulation research is the lack of confirmed regulatory interactions that can be used for benchmarking (discussed in Marbach et al., 2012 and Garcia Alonso et al., 2019), as well as the highly biased/imbalanced nature of the existing experimental evidence. In Morin et al. (2023) we used literature curated targets from low-throughput biochemical assays as positive labels to evaluate the aggregation of ChIP-seq and TR perturbation experiments, an approach we also apply in this study.

TR-target interactions supported by low-throughput experimental evidence were collected from our prior study (Chu et al., 2021), which compiled information from other resources (TRRUST: Han et al., 2018; InnateDB: Lynn et al., 2008; TFactS: Essaghir et al., 2010; TFe: Yusuf et al., 2012; HTRIdb: Bovolenta et al., 2012; CytReg: Carrasco Pro et al., 2018; ORegAnno: Lesurf et al., 2016; ENdb: Bai et al., 2020) and then significantly expanded upon neurologically-relevant TRs. Since Chu et al. (2021) was published, we have further expanded this collection, to a total of 27,629 experiments encompassing 772 TRs and 5,899 gene targets. Note that a single TR-target interaction may be supported by more than one experiment. For all analyses, we considered the presence of any experiment (e.g., EMSAs, reporter assays) in this collection supporting an interaction as a positive label. If a target gene was in the orthologous gene set, we allowed it to be counted as a label for either species.

There are several key considerations to this benchmark. First, there is an imbalance of curated targets (positive labels) for each TR, coupled with the absence of definitive negative interactions — thus all genes lacking curation are treated as negatives. It is thus a near certainty that the negative labels for any given TR contain true gene targets that have yet to be tested/curated. Second, the precision-recall (AUPRC) and receiver operating characteristic (AUROC) values were generally better than random but still modest (Fig. 3B). This reflects the incomplete nature of the literature curation corpus (De Smet and Marchal, 2010) and the complexity of benchmarking gene regulation, where no single line of regulation evidence is exhaustively performant (Garcia-Alonso et al, 2019).

We note that other works may sample the non-curated genes to be size-matched to the positive set (e.g., as in Garcia Alonso et al., 2019). We choose to avoid this balancing strategy, finding that it artificially boosts the raw AUC values. In addition, each TR has a variable count of curated targets: while this is not a problem for comparing aggregate profiles *within* a TR (i.e., ASCL1’s aggregate binding versus coexpression profiles), it complicates comparison *across* TRs. In most of our reporting, we focus on how well the observed AUC compares against null AUCs via its quantile (where a quantile of 1 means that the observed AUC was better than every null iteration). This means that each reported AUC quantile is relative to its own specific null expectation. Hence, our focus is on evaluating whether these aggregate profiles have predictive power (are better than random) at prioritizing their own curated targets to provide a relative sense of their regulation information content.

We acknowledge that one caveat of this quantile approach is that it loses information about the raw AUC value. As an example, if a TR had an AUROC of 0.67 for its aggregate coexpression profile and 0.62 for its binding profile, and both of the values were better than all null comparisons, each would achieve a quantile of 1, even though the coexpression performance was slightly higher. Still, we believe that using these quantiles (a representation of performance versus a null) is a fairer means to compare values across TRs. See Supplemental Fig. 5D for the distributions of the raw AUROCs for the aggregated positive coexpression, binding, and integrated profiles.

**Supplemental Figures**

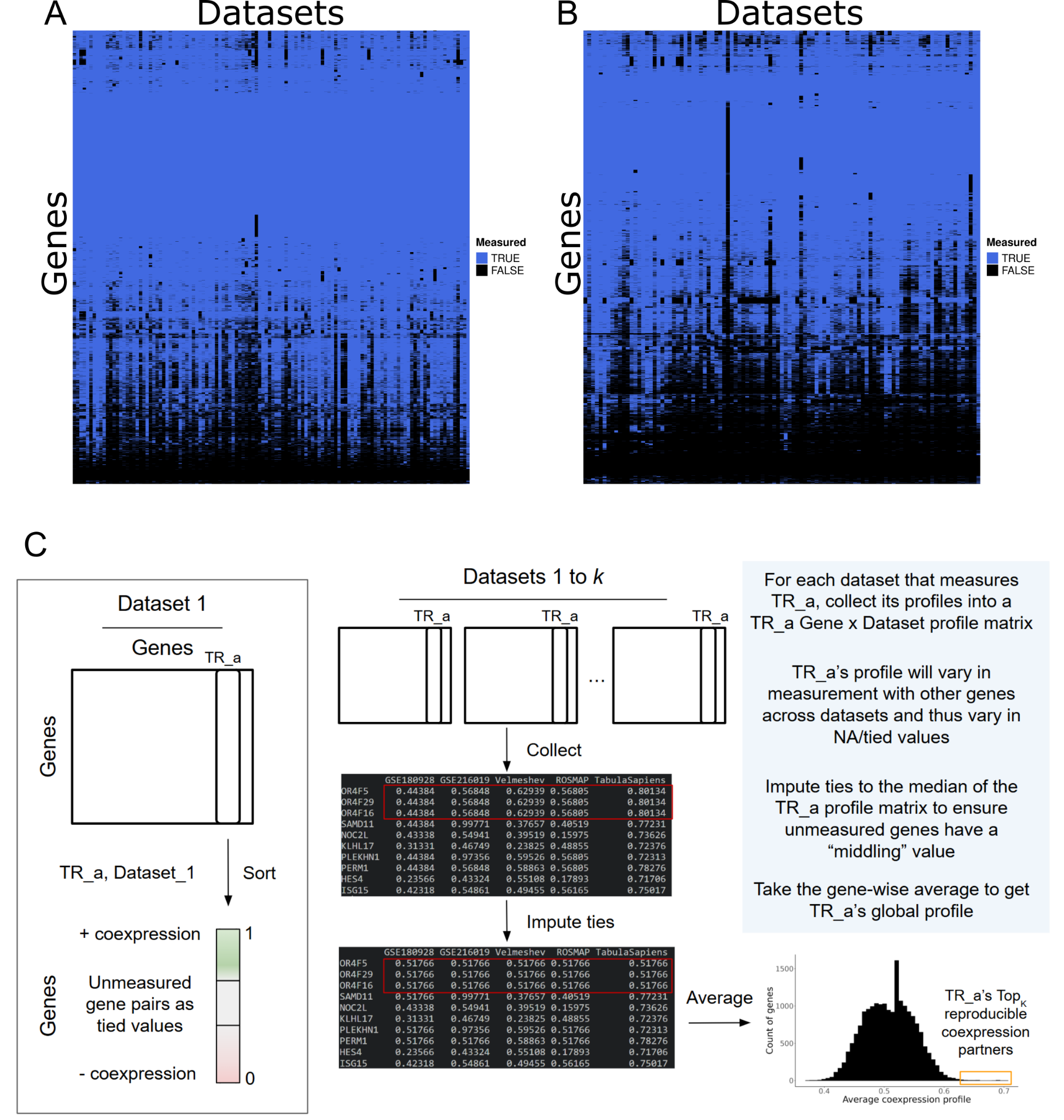

**Supplemental Figure 1.** Gene measurement coverage. (A) Binary heatmap indicating whether (blue) or not (black) a gene had non-zero counts in at least 20 cells in at least one cell type in a dataset, for 19,213 human protein coding genes and 120 datasets. (B) Mouse: 20,971 protein coding genes and 103 experiments. (C) Schematic of global TR coexpression profile aggregation. Left: Each dataset results in one gene by gene coexpression matrix by aggregating across cell types (schematized in Fig. 1C), from which a single gene coexpression profile can be extracted. Right: A given gene’s profile (e.g., “TR_a”) can be extracted from each dataset. As each profile/dataset will vary in its gene measurement, unmeasured/tied values are imputed to the median value of all of TR_a’s profiles before averaging across genes to get TR_a’s global coexpression profile.

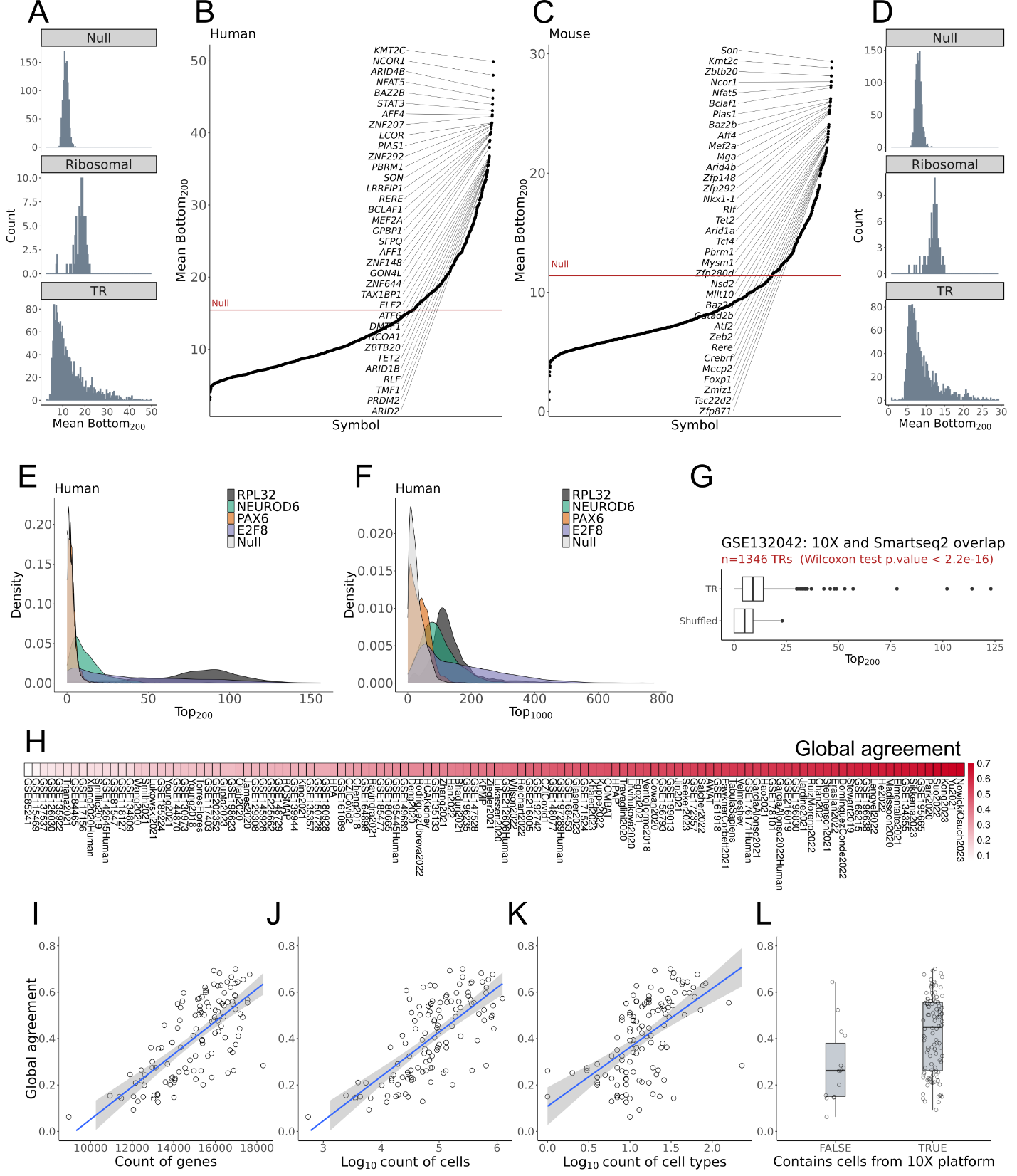

**Supplemental Figure 2.** Similarity of negative correlation TR profiles across datasets. (A) Top panel: Histogram of 1000 iterations of sampling one TR profile from each of 120 human datasets and calculating the average size of the Bottom_200_ overlap between every pair of sampled profiles, representing a null background setting. Middle panel: Histogram of the average Bottom_200_ overlap of all dataset pairs for each of 82 ribosomal genes. Bottom panel: Histogram of the average Bottom_200_ overlap of all dataset pairs for 1,605 human TRs. (B) The average Bottom_200_ overlap of all human TRs, with the red line indicating the best null overlap. (C,D) Same as in A,B, save for 103 mouse experiments and 1,484 TRs. (E,F) The distribution of (C) Top_200_ and (D) Top_1000_ overlaps between every pair of PAX6 and NEUROD6 profiles in human, with ribosomal RPL32, TR E2F8, and a representative null sample included for reference. (G) The distribution of overlaps between TR profiles generated from the same study but using different technology. (H) Each human dataset’s *Global agreement*, a measure averaging how well the TR profiles for a dataset aligned with the global TR profiles. (I-L) Plotting each dataset’s *Global agreement* against its count of (I) genes, (J) cells, (K) cell types, and (L) whether the dataset included any data using the 10X platform.

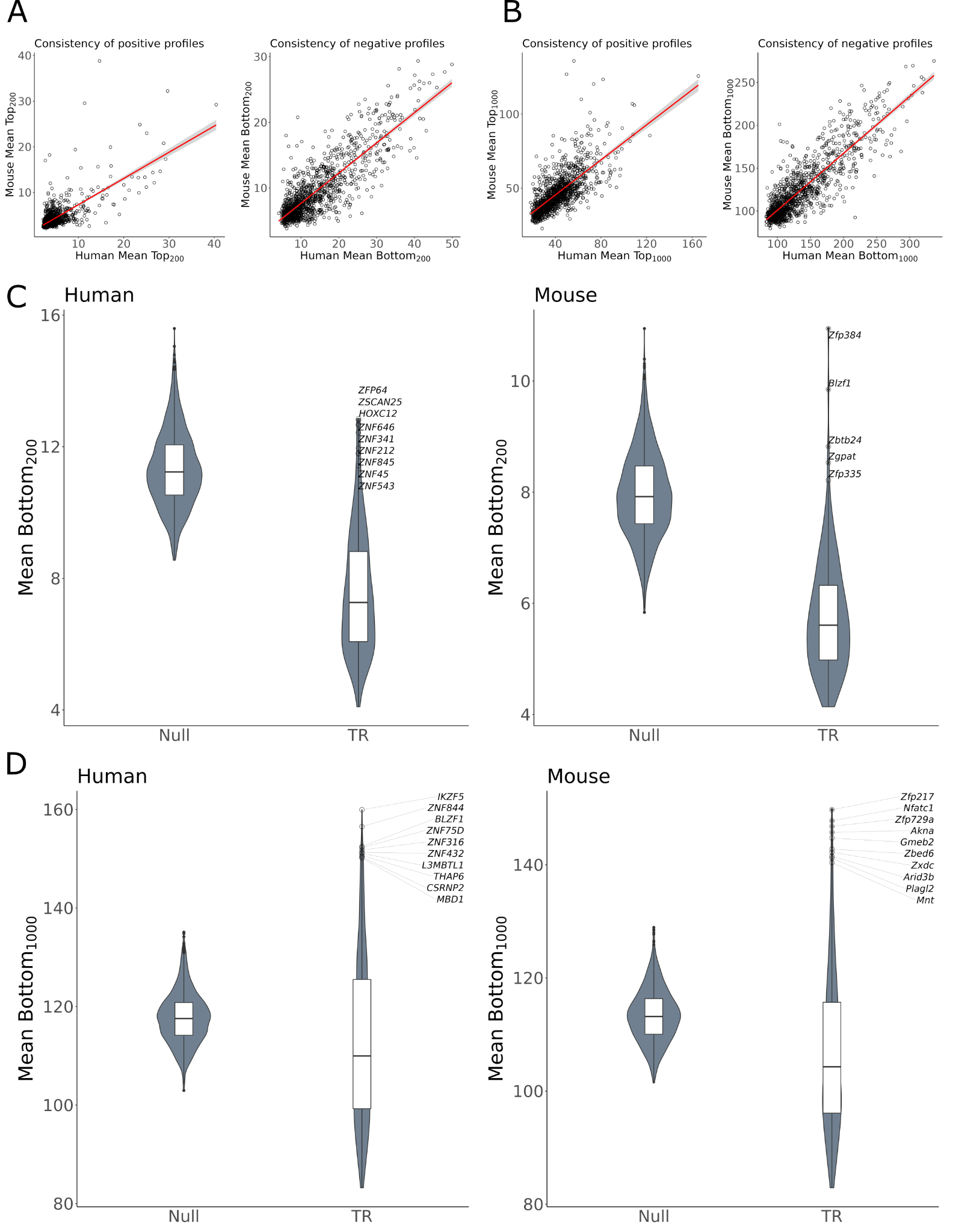

**Supplemental Figure 3.** (A, B) Conservation of the consistency of positive (Top_K_) and negative (Bottom_K_) profiles between mouse and human for 1,228 orthologous TR at (A) K=200 and (B) K=1000. Each point represents a TR with a one-to-one ortholog between mouse and human. (C, D) Examples of TR profiles with consistent negative, but not positive, profiles at (C) K=200 and (D) K=1000. The TR group considers only TRs whose Top_K_ values were lower than the typical null Top_K_ value. The Null group shows the range in mean Bottom_K_ values of all 1000 shuffled null comparisons.

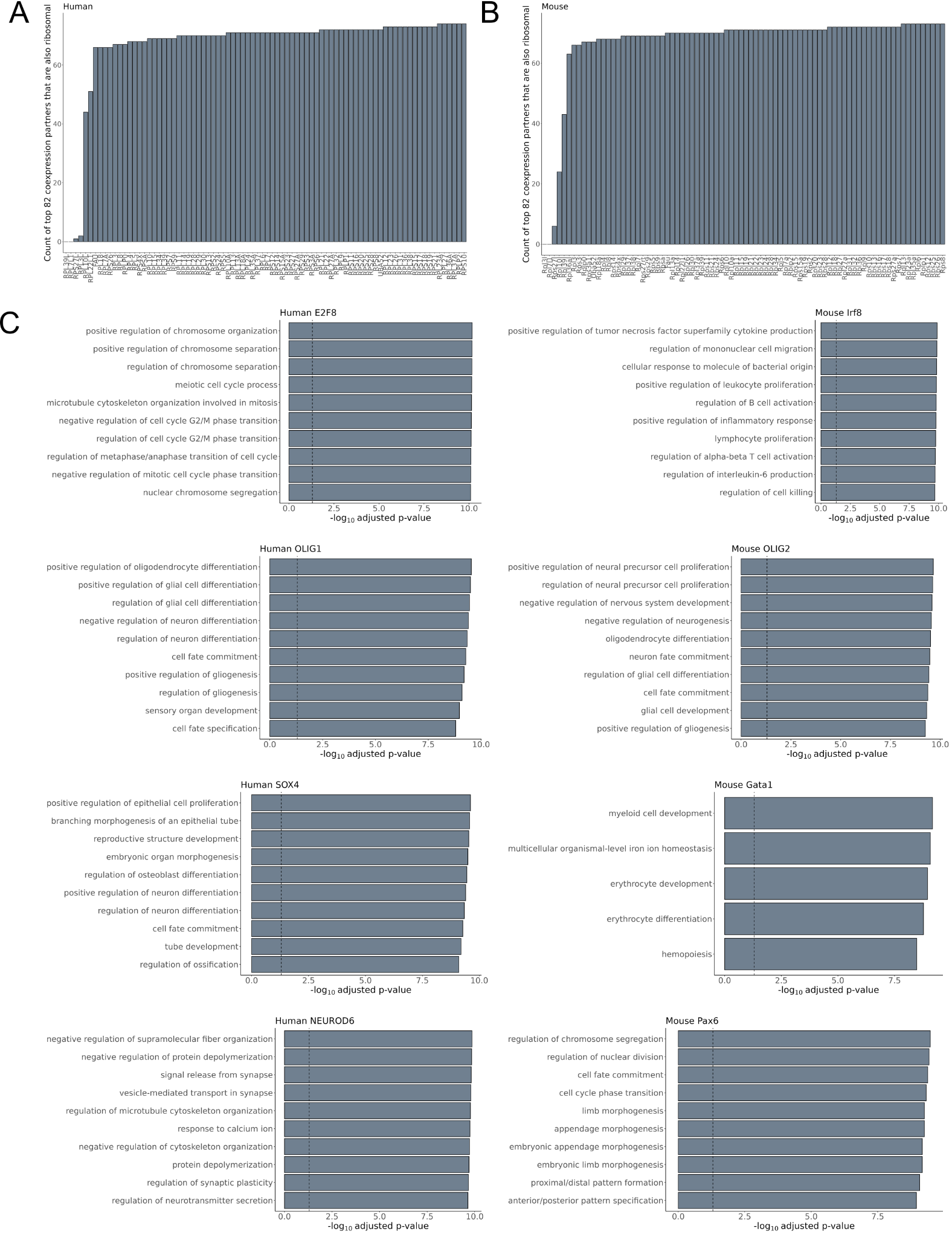

**Supplemental Figure 4.** L/S ribosomal gene aggregate profiles prioritize other ribosomal genes in (A) human and (B) mouse. Each bar represents one of the 82 L/S ribosomal genes with an unambiguous one-to-one ortholog between mouse and human. For each ribosomal gene we aggregated its coexpression profiles and then calculated how many of its top coexpressed partners belonged to the set of 82 ribosomal genes. (C) The top 10 enriched GO terms affiliated with the aggregated coexpression profiles of selected TRs.

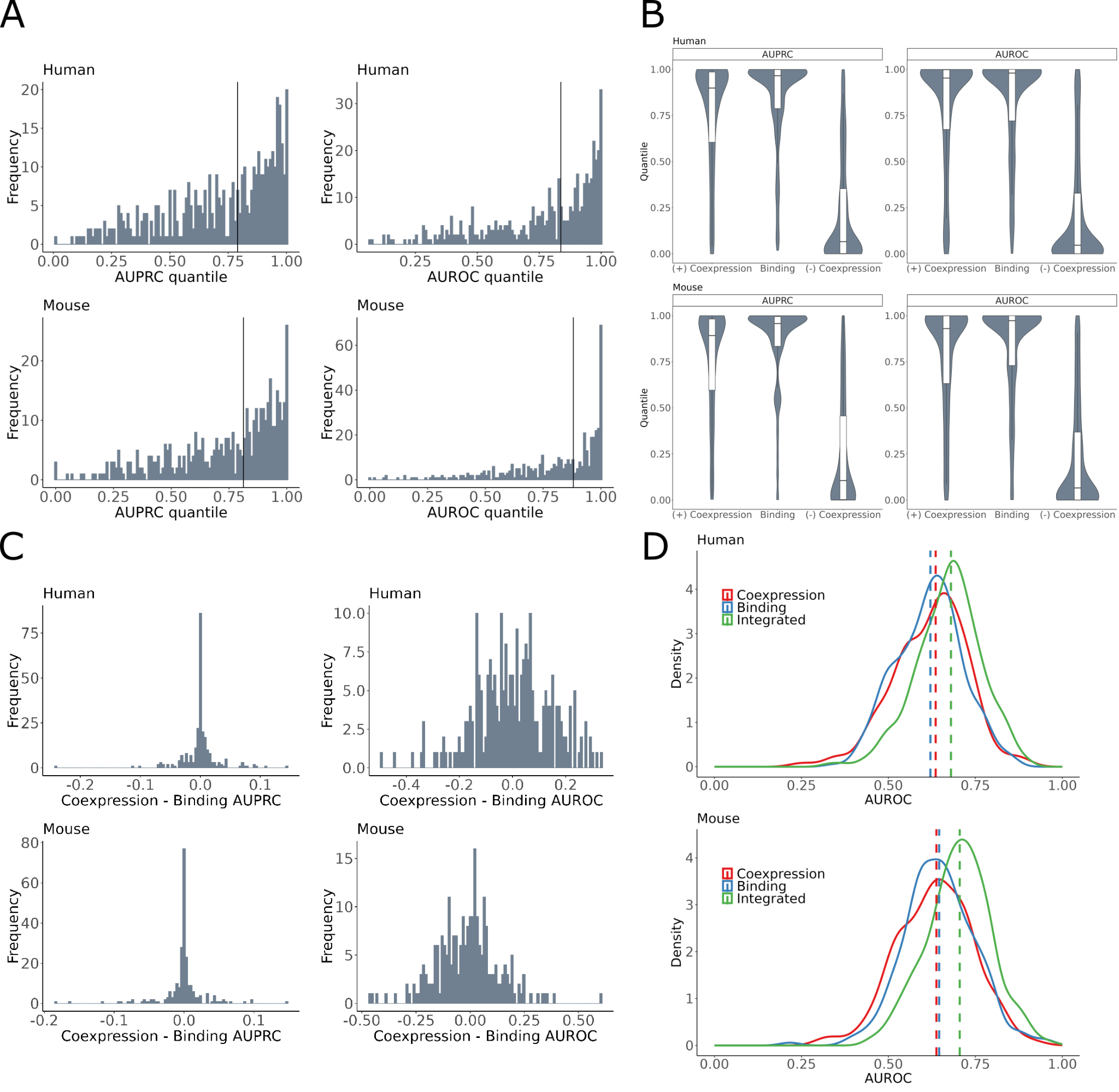

**Supplemental Figure 5.** Literature curated target benchmark. (A) Aggregating TR coexpression profiles tends to improve recovery of curated targets. Shown are the histograms of the observed AUC quantiles for 451 human and 434 mouse aggregate TR coexpression profiles. For each TR, an AUC is generated for every dataset’s ability to recover the given TR’s curated targets. This process is repeated for the TR’s aggregate profile, which is then compared to all of the individual dataset AUCs. A quantile of 1 indicates that an aggregate profile had a higher AUC (assigned better ranks to curated targets) than all of the individual dataset profiles that compose the aggregate. Black lines correspond to the median aggregate AUC quantile. (B) Distributions of the AUC quantiles for aggregated negative/positive coexpression and binding profiles for the 253 human and 241 mouse TRs that had binding and coexpression data. (C) Histograms of the difference between the raw AUC values between coexpression and binding aggregates. Positive values indicate that coexpression was better able to recover curated targets, negative values indicate binding data was better. (D) Integrating the positive coexpression and binding aggregates (via rank product) tends to increase recovery of curated targets.

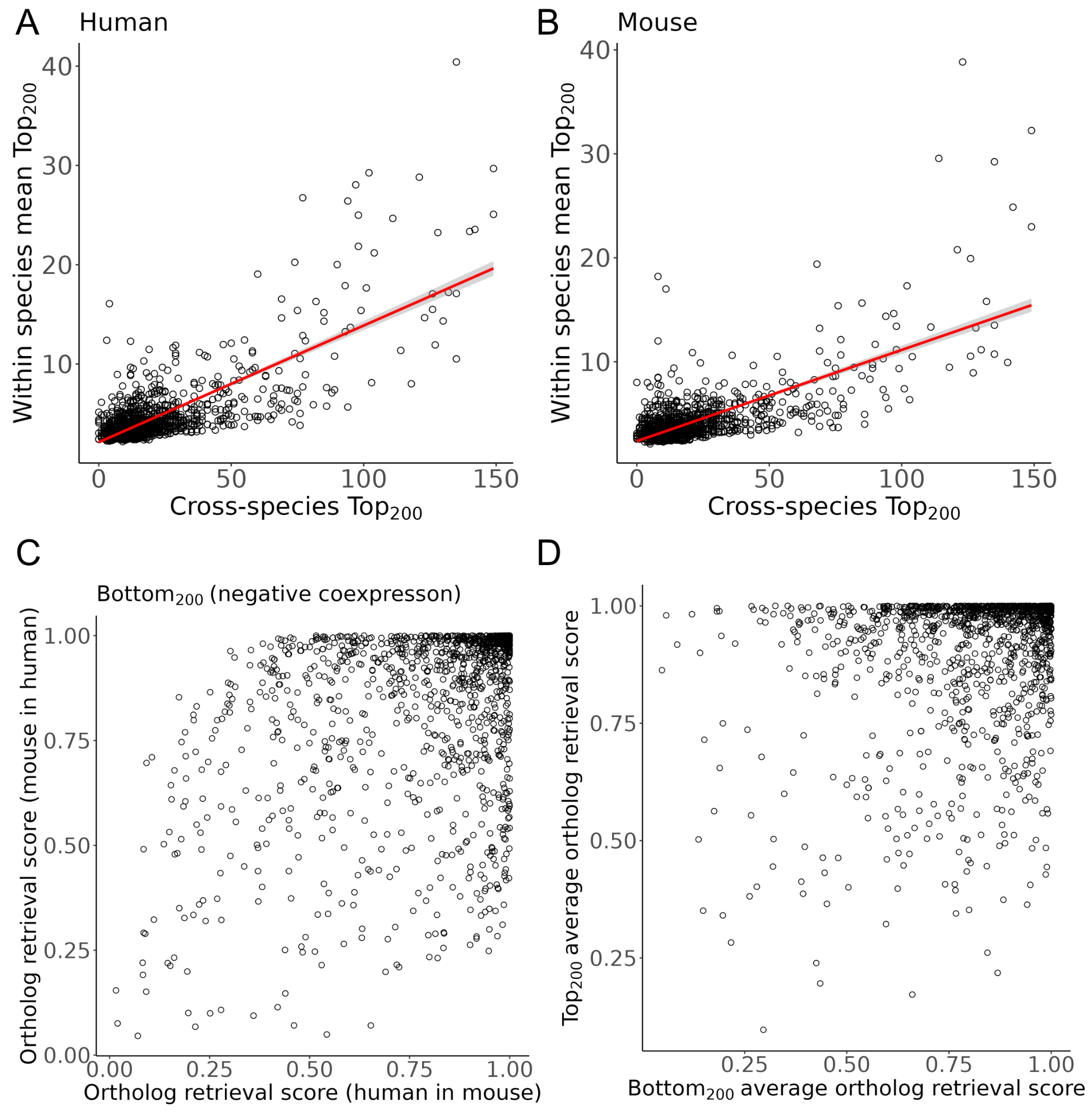

**Supplemental Figure 6.** Comparison of single cell coexpression across species. (A, B) TRs that are consistent within species tend to be consistent across species. Scatterplots of the *Top_200_* overlap between orthologous TRs versus the average *Top_200_* between every unique pair of individual TR profiles in (A) human and (B) mouse. Each point is an orthologous TR, the y-axis is a measure of TR coexpression agreement before aggregation within each species, while the x-axis is common to A and B and represents the agreement between aggregate ortholog profiles between species. (C) Scatterplot of the *Bottom_200_* ortholog retrieval scores. (D) Scatterplot of the averaged *Bottom_200_* and *Top_200_* ortholog retrieval scores: the upper right quadrant indicates TRs whose positive and negative coexpression is consistent and specific between species.
**Supplemental Citations**
